# CRISPR/Cas9 mediated mutations as a new tool for studying taste in honeybees

**DOI:** 10.1101/2020.03.26.009696

**Authors:** Laura Değirmenci, Dietmar Geiger, Fábio Luiz Rogé Ferreira, Alexander Keller, Beate Krischke, Martin Beye, Ingolf Steffan-Dewenter, Ricarda Scheiner

**Author notes:** Corresponding author: University of Würzburg, Behavioral Physiology and Sociobiology, Biocenter, Am Hubland, 97074 Würzburg, Germany.

## Abstract

**Background:** Honeybees rely on nectar as their main source of carbohydrates [1]. Sucrose, glucose and fructose are the main components of plant nectars [2] [3]. Intriguingly, honeybees express only three putative sugar receptors (AmGr1, AmGr2 and AmGr3) [4], which is in stark contrast to many other insects and vertebrates. The sugar receptors are only partially characterized [5] [6]. AmGr1 detects different sugars including sucrose and glucose. AmGr2 is assumed to act as a co-receptor only, while AmGr3 is assumedly a fructose receptor.

**Results:** We show that honeybee gustatory receptor AmGr3 is highly specialized for fructose perception when expressed in *Xenopus* oocytes. When we introduced nonsense mutations to the respective *AmGr3* gene using CRISPR/Cas9 in eggs of female workers, the resulting mutants displayed almost a complete loss of responsiveness to fructose. In contrast, responses to sucrose were normal. Nonsense mutations introduced by CRISPR/Cas9 in honeybees can thus induce a measurable behavioural change and serve to characterize the function of taste receptors *in vivo*.

**Conclusion:** CRISPR/Cas9 is an excellent novel tool for characterizing honeybee taste receptors *in vivo*. Biophysical receptor characterisation in *Xenopus* oocytes and nonsense mutation of *AmGr3* in honeybees unequivocally demonstrate that this receptor is highly specific for fructose.

**Graphical Abstract:** Figure 0

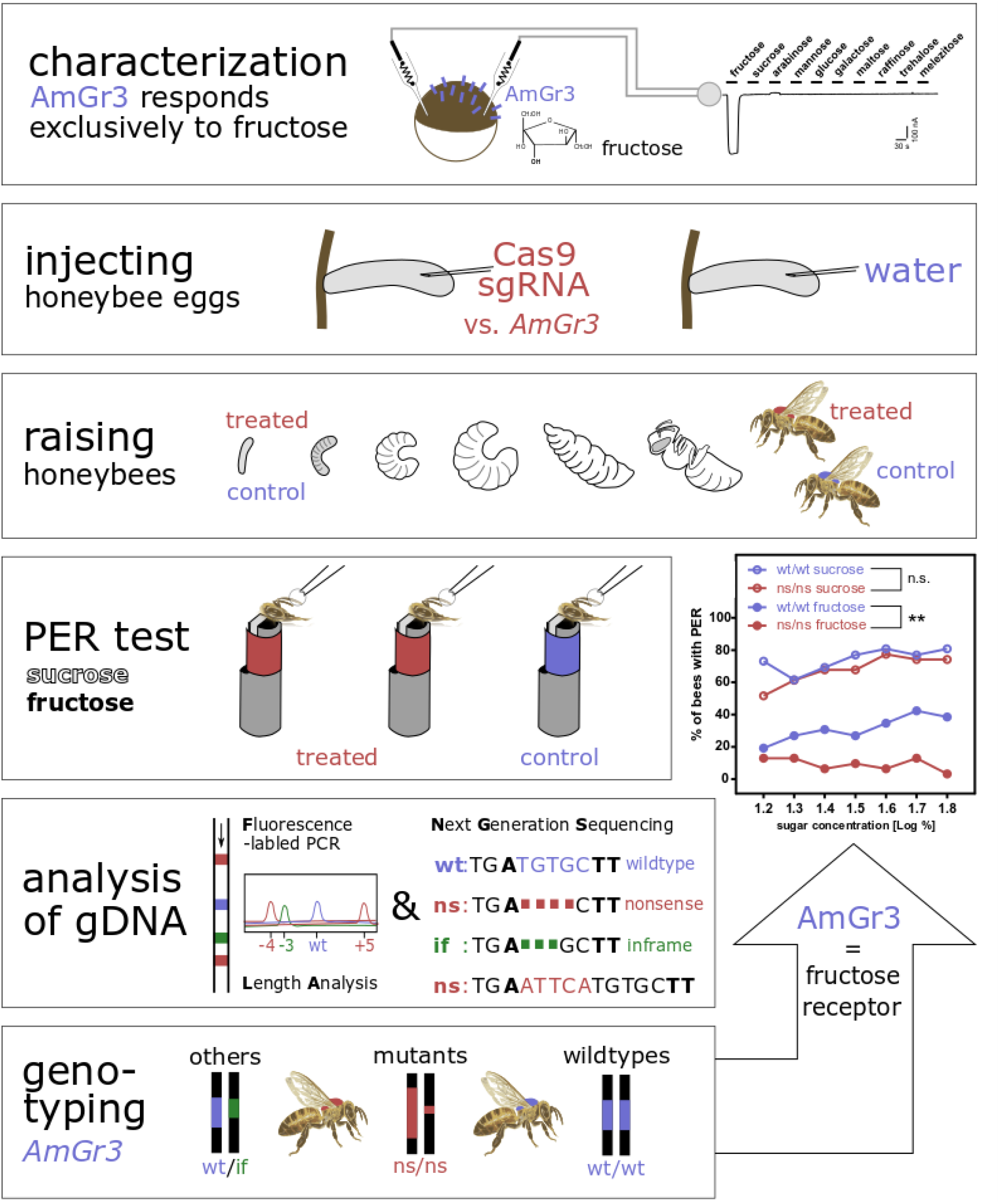

## Background

Honeybees (*Apis mellifera*) are not only important pollinators world-wide. The highly social insects perform an intricate division of labour and are well-known for their astonishing skills in learning and communication. When it comes to taste, however, honeybees display a rather poor set of receptors. Because plant-derived nectar is their sole source of carbohydrates, sugar perception is naturally of utmost importance for honeybees. The bees sense the sugar composition of a food source with only a few fine contact chemoreceptors on their antennal tip [7]. In contrast to many other insects such as the fruit fly (*Drosophila melanogaster*) with 68 genes and mosquitoes (*Anopheles gambiae*) with 75 genes, the genome of the honeybee comprises only ten genes coding for gustatory receptors (Grs) [4]. Among these only three code for sugar receptors: AmGr1, AmGr2 and AmGr3 [4] [8]. The taste receptors are expressed in the brain, the antennae, mouthparts, tarsi and the gut of the honeybee. With this small set of receptors honeybees evaluate a diverse set of sugars like sucrose, fructose, maltose and melicitose in nectar in varying composition and in amounts ranging from 5% to 80%. In flowers of mint plants (*Laminacea*), buttercups and clematis (*Ranunculaceae*), for example, sucrose is the main sugar, whereas other flowers such as those of oilseed rape contain relatively more glucose and fructose [3] [9]. The sugar trehalose, in contrast, acts as blood sugar [10] [2].

How honeybees recognize the different sugars in nectar and in the inner organs with this small set of receptors is unclear. While AmGr1 was shown to detect a variety of sugars (sucrose, fructose, glucose and trehalose), AmGr2 seems to function as co-receptor only [5]. AmGr3 appears to specifically perceive fructose [6], but not all of the relevant sugars have been tested so far. The AmGr3 receptor is an ortholog of the *Drosophila* fructose receptor DmGr43a [4] and is similarly affine for fructose as the BmGr9 of the silkworm *Bombyx mori* [11]. Because AmGr3 appears to selectively respond to one sugar, it is an interesting candidate for characterizing its function through a nonsense-mutation in the *AmGr3* gene.

The function of insect and mouse taste receptors has been frequently characterized using electrophysiological techniques with heterologously expressed receptors in frog oocytes (*Xenopus* oocytes, [5] [6]), human liver cells (HEK cells, [11]) or plant cells (*Arabidopsis* mesophyll protoplasts, [12]). However, experimental indications from heterologous expression systems need to be verified in the original organism using knock out or knock down mutants, such as has frequently been performed in fruit flies (for review see [13]).

While techniques of genetic manipulation are generally not very successful in honeybees, the CRISPR/Cas9 system is a promising new genome-editing technique which has been employed successfully in numerous insects such as *Drosophila melanogaster* and *Aedes aeqypti* ([14] and [15], respectively). Applications in honeybees are still rare. Kohno et al. (2016) managed to produce mosaic queens and mutated drones lacking a major royal jelly protein (*mrjp1* gene) [16]. Roth et al. (2019) applied CRISPR/Cas9 on honeybee workers to investigate the sex termination pathway and to identify genes influencing size polymorphism [17].

We characterized the function of the putative fructose receptor AmGr3 classically by heterologous expression in *Xenopus laevis* oocytes and elucidated its cation transport characteristics through two-electrode voltage-clamp technique (TEVC). In addition, we employed the novel CRISPR/Cas9 technique to induce specific mutations of this receptor in live honeybees and tested their responsiveness to different sugars as one-week old adults.

## Results

### AmGr3 represents a hyperpolarisation-activated fructose receptor

Our results demonstrate that AmGr3 is clearly a fructose receptor. Upon addition of 160 mM fructose to the external solution, AmGr3-expressing oocytes elicited inward cation currents (negative currents) with amplitudes of several hundred nano amps at a holding potential of −80 mV (Fig. 1A). Removing the fructose from the bath medium, the inward currents returned to the pre-fructose level. Control oocytes did not show any fructose-induced currents (Fig.1A lower panel). To study the voltage dependence of AmGr3 mediated currents, 200 ms test voltage pulses were applied in the range from +10 mV to −150 mV in 20 mV decrements in absence and presence of fructose (Fig. 1B). Fructose-induced currents were derived by subtracting the currents in the absence of fructose from the currents in its presence (Fig. 1B and C). The derived fructose-induced currents are characterized by time-dependent activation kinetics (Fig. 1B) and hyperpolarization-dependent activation (Fig. 1C).

**Fig. 1:**
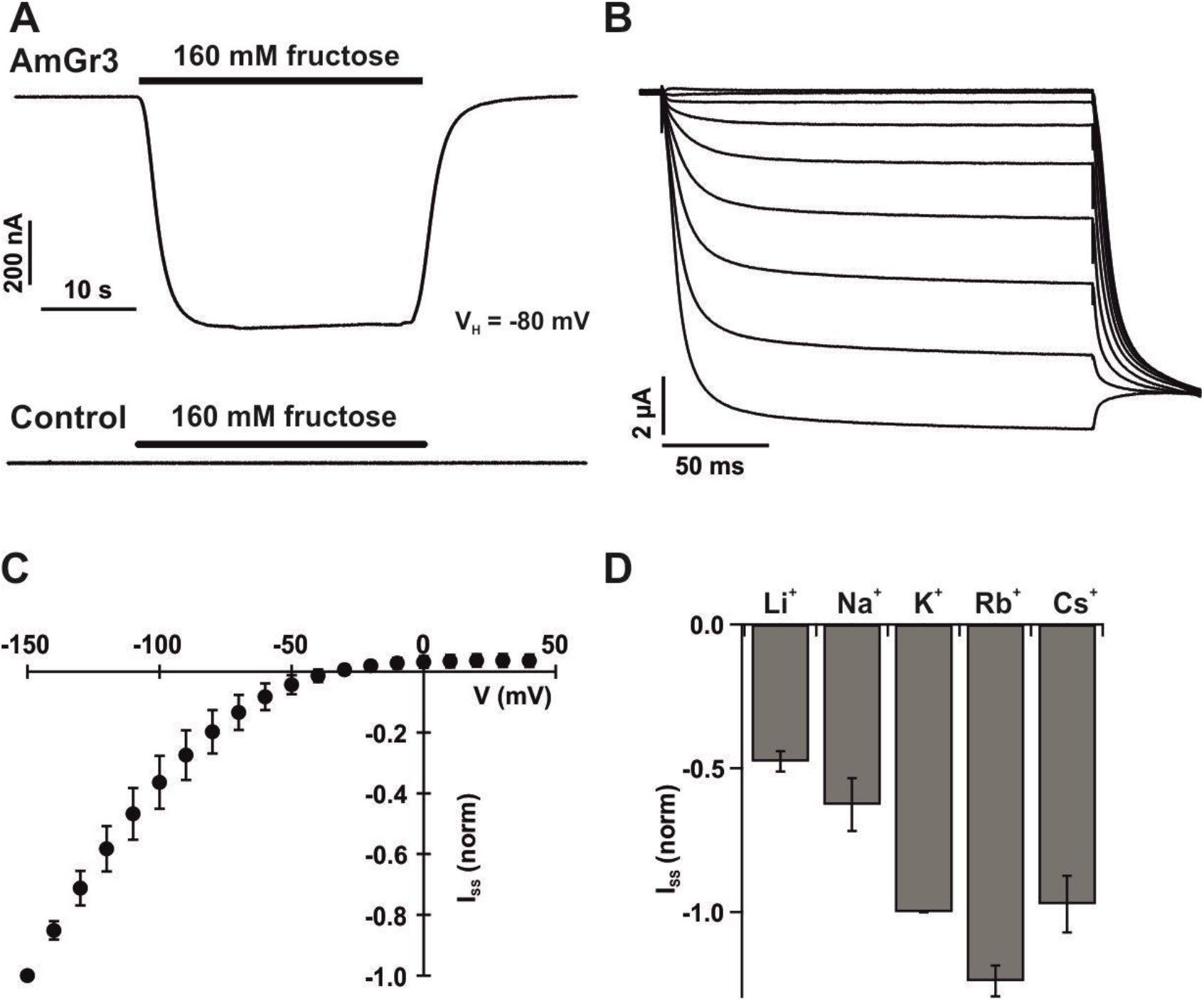
AmGr3 represents a hyperpolarisation activated fructose receptor. **A)** Representative whole oocyte currents recorded at −80 mV in response to perfusion with fructose in standard solution. Upper panel: AmGr3 expressing oocyte; Lower panel: non-injected control oocyte. **B)** Representative fructose-induced whole oocyte currents in response to a series of 200 ms test pulses ranging from +10 mV to −150 mV in 20 mV decrements. Each test pulse was followed by a constant voltage pulse to −140 mV. The holding potential was at 0 mV. Currents were recorded in standard solution containing 160 mM fructose. **C)** Fructose-induced steadystate currents (I_SS_) from AmGr3 expressing oocytes were plotted as a function of the applied membrane potential. Fructose-induced currents were derived by subtracting the currents recorded in standard solution containing 160 mM sorbitol from the currents in standard solution containing 160 mM fructose (n = 3 ± SD). **D)** Fructose-induced I_SS_ were recorded in the presence of 30 mM of different monovalent cations (as indicated) and 160 mM fructose at a membrane potential of −140 mV. AmGr3 derived currents were normalized to the currents in K^+^-based media (mean of n = 5 oocytes ± SE).

### Fructose-activated AmGr3 mediates non-selective cation currents

Gustatory receptors represent a group of (non-GPCR) seven-transmembrane receptors that detect tastants (non-volatile compounds) via contact chemo sensation. Upon ligand binding, these receptors elicit cation currents finally leading to the firing of action potentials in gustatory neurons [11]. To test the selectivity of the receptor for cations, oocytes expressing the gustatory receptor AmGr3 were perfused with external solutions containing 30 mM of different monovalent cations. The fructose-induced ionic currents were recorded at a membrane potential of −140 mV. In response to fructose perfusion, negative current deflections appeared in all cationic conditions tested (Fig. 1D). To calculate the relative permeability of AmGr3 for cations, reversal potentials in the presence of different cations and fructose were monitored. Reversal potentials appeared similar between the cations tested. AmGr3 thus seems to be a rather non-selective cation channel with a relative permeability sequence of K^+^=1±0 > Rb^+^ = 0.97 ± 0.05 > Cs^+^ = 0.91 ± 0.05 > Na^+^ = 0.83 ± 0.09 > Li^+^ = 0.70 ± 0.06 (permeability of K^+^ was set to 1, mean of n = 5 oocytes ± SE).

### Fructose is the only sugar inducing AmGr3-derived currents

In 2018, Takada et al. reported that the gustatory receptor AmGr3 responds only to fructose when transiently expressed in *Xenopus* oocytes [6]. To confirm these results and to broaden the list of sugars tested (by additional use of arabinose, raffinose and melezitose), we successively perfused AmGr3-expressing oocytes with different mono-, di-, and trisaccharides (160 mM each) at a membrane potential of −80 mV (Fig. 2A). Among the ten sugars tested, AmGr3 only responded to fructose, suggesting that AmGr3 is indeed a fructose specific receptor (Fig. 2A and B; cf., [6]).

**Fig. 2:**
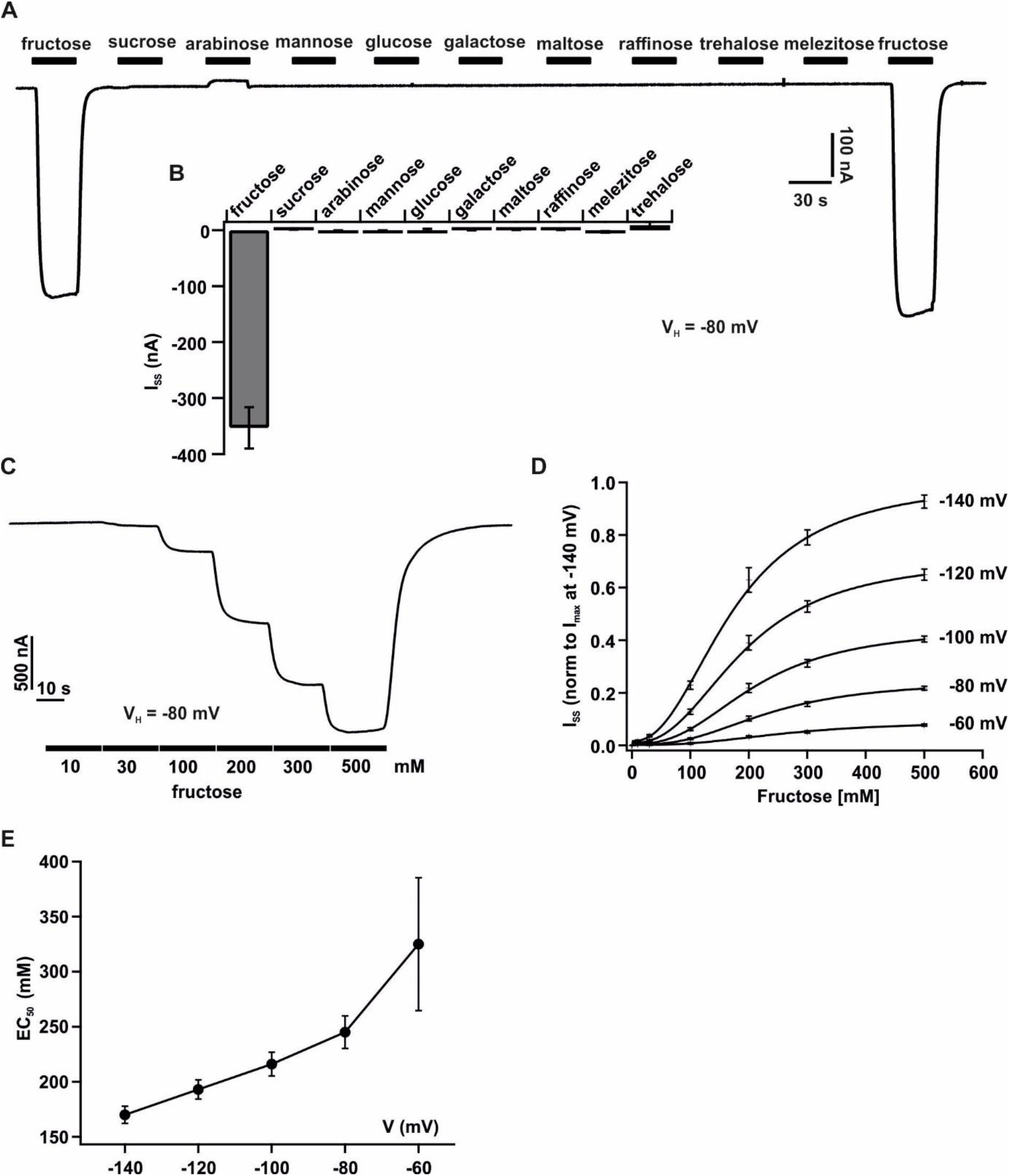
Fructose is the only sugar inducing AmGr3-derived currents. **A)** Representative whole oocyte currents from oocytes expressing AmGr3 were recorded at −80 mV in response to perfusion with 160 mM of different mono-, di- and trisaccharides in standard solution. Perfusion with test sugars are indicated by black bars. **B)** Statistical analysis of the sugar selectivity of AmGr3 expressed in *Xenopus* oocytes. Steady state currents in the presence of the indicated sugars were monitored at a membrane potential of −80 mV (mean of n = 9 oocytes ± SE). **C)** Whole oocyte current recording from an AmGr3 expressing oocyte at a membrane potential of −80 mV. Successive elevation of the fructose concentration (black bars indicate the applied fructose concentration) in the standard solution gradually increased the AmGr3-mediated currents. **D)** I_SS_ from AmGr3 injected oocytes were recorded in presence of rising extracellular fructose concentrations and plotted as a function of the fructose concentration. A Hill function was fitted to the individual fructose saturation curves at the indicated membrane potentials (black solid line; mean of n = 11 oocytes ± SE). **E)** The apparent affinity constants EC_50_ derived from fits such as shown in D) were plotted as a function of the membrane potential (mean of n = 11 oocytes ± SE).

Stepwise increases in fructose concentrations resulted in a gradual rise in AmGr3-mediated currents at a membrane potential of −80 mV (Fig. 2C). When the steady-state currents, recorded in presence of rising extracellular fructose concentrations (3 up to 500 mM), were plotted as a function of the fructose concentration, AmGr3 currents increased upon membrane hyperpolarization and started to saturate between 300 and 500 mM fructose (Fig. 2D). A Hill function sufficiently described the individual fructose saturation curves at the given membrane potentials between −60 and −140 mV (Fig. 2D). The apparent affinity constant EC_50_ of AmGr3 was 210 mM at −100 mV. Plotting the calculated EC_50_ values as a function of the membrane potential (Fig. 2E), it becomes apparent that hyperpolarizing voltages increased the apparent affinity of AmGr3 from 325 ± 60.4 mM at −60 mV to 170 ± 7.8 mM at −140 mV.

Thus, our data show that AmGr3 is indeed a highly selective fructose receptor when expressed in *Xenopus* oocytes, leading to the question whether a nonsense mutation of this gene in live honeybees could affect their behavioural response to fructose.

### CRISPR/Cas9 confirms AmGr3 as a specific fructose receptor in live honeybees

We used CRISPR/Cas9 to introduce indels (insertions or deletions) which are not a multiple of three, leading to non-functional proteins of AmGr3 [18] (for sgRNA target-site and the location of the introduced frame shift in relation to the entire ORF, exons and introns of *AmGr3*, see Fig. 3). Two replicate experiments were performed, the second with a reduced sample size due to the extreme hot and dry summer 2018 (honeybee queens adapt their egg laying performance to the nectar flow and robustness during *in-vitro* rearing decreases with low humidity). Around 80 % of all eggs injected tolerated the injection. The treatment showed a 9.6 % hatching rate (13.9 % in replicate B). Between 53.3 and 69.9 % of the control bees hatched into larvae (Tab. 1). The survival to adult emergence varied from day to day (59-86 %), likely due to the manual transferring steps (after hatching on food, before pupation on filter paper, not shown in the table). All one-week old adult bees were tested for their responses to fructose and sucrose. Only double nonsense (ns/ns) mutants and wildtypes (wt/wt) were included in the evaluation.

**Fig. 3:**
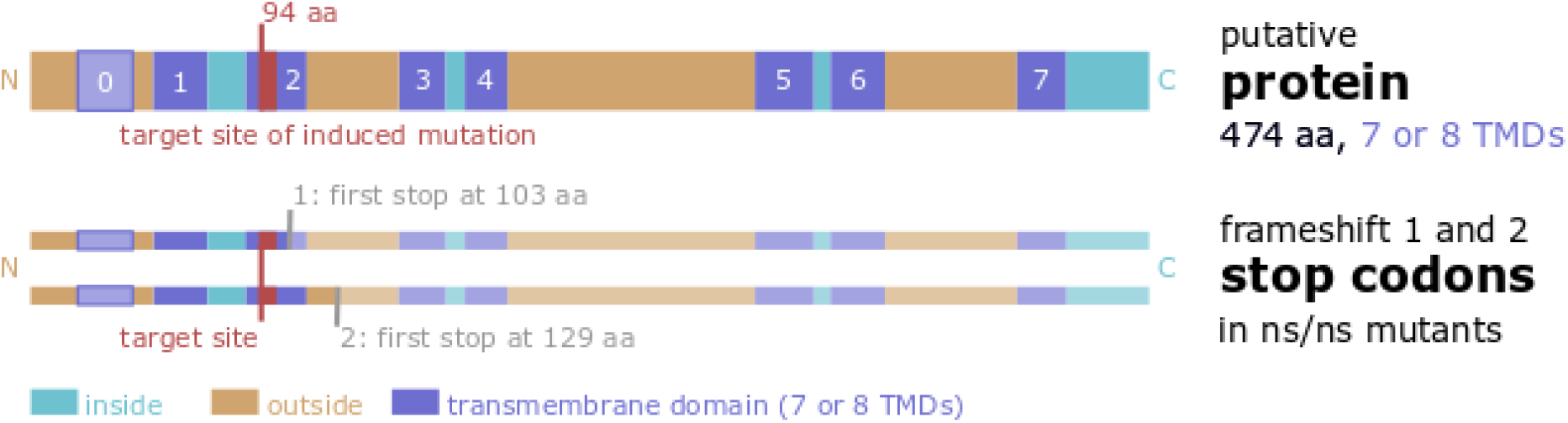
CRISPR/Cas9 induced nucleotide changes at the *AmGr3* gene target the second putative transmembrane domain and introduce double-nonsense mutations (ns/ns) at high frequency. The graph shows that the mRNA target-site for *AmGr3* (5’-gcaacttgtagtgatgtgcttgg-3’) is placed within the putative second transmembrane domain (TMD) after the N-terminus. Folding predictions (I-TASSER, PHYRE-Protein and TMHMM) show different outcomes about a possible upstream TMD (TMD 0). The two possible frameshift mutations (not a multiple of three) driven from the sgRNA target-site introduce either a stop codon at position 103 aa or 129 aa (amino acids) of the deduced sequence and are followed by multiple stops. As a consequence, five TMDs of the AmGr3 proteins are lacking in double-nonsense (ns/ns) mutants so it is assumed not to function as a fructose receptor at all.

**Table 1:** Survival and hatching numbers and rates of the sgRNA and Cas9 injected and control honeybee eggs or eggs with no injection under artificial rearing conditions. The 24h rate shows the percentage of eggs that tolerated the injection and were still intact after 24h. The hatching rate shows the percentage of larvae that hatched form the surviving eggs. The frequencies of 24h survival differs not when injected with sgRNA6 and Cas9 nuclease or water only (% 24h survival; replicate A: n.s., p=0.7970; replicate B: n.s., p=0.0895; Fisher’s exact test). The survival of microinjected eggs of 24h is statistically different from eggs, that were not injected (% 24h survival; total: ***, p<0.0001, Chi-Square-test). The hatching rate after three days decreases statistically when injected with water (% hatched; replicate A: ***, p<0.0001; replicate B: *, p=0.0126; Fisher’s exact test) and even lower when injected with sgRNA and Cas9 (% hatched; total: ***, p<0.0001; Chi-Square test).

|  | replicate A |  | replicate B |  | total |  |
| --- | --- | --- | --- | --- | --- | --- |
| treatment: | # 24h | # hatched | # 24h | # hatched | # 24h | # hatched |
| sgRNA and Cas9 injected | 1,436 | 200 | 1,116 | 107 | 2,552 | 307 |
| water injected | 90 | 48 | 94 | 65 | 184 | 113 |
| no injection | - | - | - | - | 251 | 183 |
| treatment: | % 24h | % hatched | % 24h | % hatched | % 24h | % hatched |
| sgRNA and Cas9 injected | 82.0 | 13.9* | 78.3 | 9.6* | 80.3 | 12.0** |
| water injected | 83.3 | 53.3 | 85.5 | 69.1 | 84.4 | 61.4* |
| no injection | - | - | - | - | 96.9*** | 72.9 |

Double nonsense mutations of the putative fructose receptor AmGr3 were not lethal during larval development and the first week of adult life. This indicates that *AmGr3* is not essential for life-preserving behaviours such as food intake. To pre-screen the effectiveness of our treatment, we performed a fluorescence length analysis (FLA) based on capillary gel electrophoresis with HEX-labelled PCR products of the bees. With FLA we detected 36.0 % double-nonsense mutants in the treatment group (49.1 % in replicate B) and 91.7 % wild types (8.3 % still with one wt and one ns allele) in the control group (100 % in replicate B). We subsequently sequenced the respective amplicons of all primal genotyped mutants (ns/ns) and wild types (wt/wt) using next generation sequencing (NGS). Our results include all individuals with assured wildtype or mutant genotype via deep sequencing of the target amplicons (NGS proofed 85 samples (86.7 %) of the FLA pre-screened genotypes). All other genotypes (allele combinations of wt, ns and if (in-frame) were disregarded, since a clear statement about the presence and functionality of their AmGr3 proteins and the measured behaviour is not possible.

In both replicates double mutants (ns/ns in fructose receptor gene *AmGr3*) displayed a significantly reduced responsiveness to fructose, unlike wildtypes (wt/wt) (Fig. 4, statistics also in Tab. 2) when tested at their antennae with rising sugar concentration [19]. Responses to sucrose, in contrast, were unaffected in both groups (Fig. 4, statistics also in Tab. 2).

**Fig. 4:**
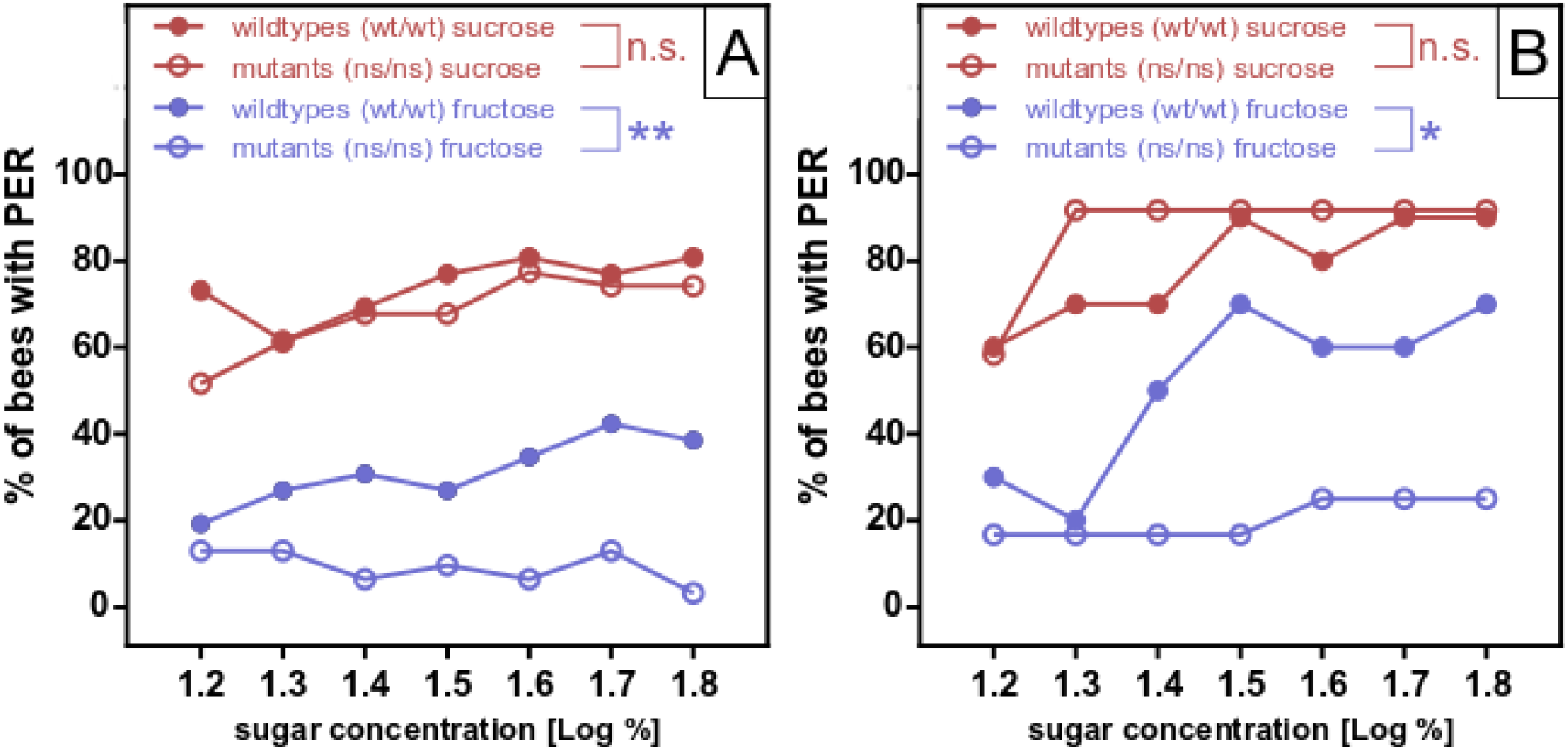
*AmGr3* mutants display a reduced responsiveness to fructose but not to sucrose. The figures (of replicate A and replicate B) show the percentage of bees responding to a defined sugar concentration of either fructose or sucrose (16 %, 20 %, 25 %, 32 %, 40 %, 50 % and 63 %, corresponding to a log of 1.2, 1.3, 1.4, 1.5, 1.6, 1.7 and 1.8). Honeybee mutants of the fructose receptor *AmGr3* gene (ns/ns - double mutants, circles) are less sensitive to increasing fructose concentrations (black) than wildtype bees (wt/wt, dots) (logistic regression [factor genotype] for **fructose - A:** **p=0.005, X^2^_1,399_=8.026, N_(wt/wt)_=26, N_(ns/ns)_=31 and **fructose - B:** *p=0.022, X^2^_1,154_=5.265, N_(wt/wt)_=10, N_(ns/ns)_=12). The same groups do not differ in their sucrose responsiveness (grey) (logistic regression [factor genotype] for **sucrose - A:** n.s. p=0.502, X^2^_1,399_=0.451, N_(wt/wt)_=26, N_(ns/ns)_=31 and **sucrose - B:** n.s. p=0.446, X^2^_1,154_=0.504, N_(wt/wt)_=10, N_(ns/ns)_=12). For statistics also see Table 2.

**Table 2:** Statistical values of logistic regression for Fig. 4 displaying that wildtype (wt/wt) and AmGr3 mutant bees (ns/ns, double nonsense) differ statistically in their response to fructose but not to sucrose. Logistic regression was performed with the factor genotype for comparing the sugar curves (fructose OR sucrose) of both treatment groups (ns/ns vs. wt/wt). Mutant bees show a reduced fructose responsiveness but both groups do not differ in their sucrose response. For graphical display see Fig. 4.

| compare sugar response curves (wt/wt vs. ns/ns of fructose OR sucrose) – logarithmic regression |  |  |  |  |  |  |  |  |  |  |
| --- | --- | --- | --- | --- | --- | --- | --- | --- | --- | --- |
| see Fig. 4 | test groups (N) |  | log. Regression |  | fructose responsiveness |  |  | sucrose responsiveness |  |  |
|  | wt/wt | ns/ns | df | N | Sign. | P | X<sup>2</sup> | Sign. | p | X<sup>2</sup> |
| replicate A | 26 | 31 | 1 | 399 | ** | 0.005 | 8.026 | n.s. | 0.502 | 0.451 |
| replicate B | 10 | 12 | 1 | 154 | * | 0.022 | 5.265 | n.s. | 0.446 | 0.504 |

## Discussion

Although honeybees rely on feeding nectar, their genome only encodes for a small set of gustatory receptors. The behavioural repertoire linked to a limited number of sugar resources [20] [21] [19] [22] (for review see [23]) and the low number of taste receptors predestine the honeybee as an interesting organism to investigate the mechanisms of taste perception. Of the ten putative honeybee taste receptors, AmGr1 (AmGr2 as its possible co-receptor) was characterized as a sugar receptor for various ligands [5] while AmGr3 is regarded as a conserved ortholog of the fructose receptor of flies and moths [24] [11] [6].

Here we reconstituted the responses of AmGr3 to various sugars in the heterologous expression system of *Xenopus* oocytes. Two-electrode voltageclamp studies revealed a fructose-specific nonselective cation current conductance in AmGr3-expressing oocytes [6]. Although genetic studies using *Drosophila melanogaster* suggest that co-expression of multiple Grs is necessary for sugar perception [25] [26], AmGr3 did not require the co-expression of other Gr subunits to respond to fructose in oocytes, just like BmGr9 and DmGr43a [11]. The broad unspecific cation conductance we found is well in line with the studies of the AmGr3 ortholog from silkworm (*Bombyx mori* Gr9, BmGr9; [11]). Sato et al. (2011) demonstrated that BmGr9 constitutes a ligand-gated non-selective cation channel [11]. Just like BmGr9, AmGr3 conducted all monovalent cations tested (Fig. 1D). Very recently, a Cryo-EM-derived three-dimensional structure of the olfactory receptor Orco (Odorant receptor co-receptor) from the parasitic fig wasp *Apocrypta bakeri* was resolved at 3.5 Å resolution [27]. The 3D structure shows that the functional receptor consists of four monomers symmetrically arranged around a central ion-conducting pore. Since Ors and Grs share the same gene structure, a predicted topology with seven transmembrane domains and a conserved motif in TM7 [28], it is tempting to speculate that functional Grs consist of four subunits, too. Whether AmGr3 assembles to heterotetrametric receptors with other members of the honeybee Gr-family and how this might influence the ligand specificity of the receptors remains to be shown. In future, the 3D structure of AbOrco will guide structure-function research not only of Ors but also of the related Gr family members.

Interestingly, fructose-induced cation currents across AmGr3 appeared activated by hyperpolarisation and thus inward rectifying (Fig. 1C; cf. [11]). Moreover, AmGr3 showed a time-dependent activation kinetics at hyperpolarized membrane potentials (Fig. 1B) and voltage-dependent EC_50_ values for fructose (Fig. 2D and E). These voltage-dependent electrical characteristics of AmGr3 require the presence of a voltage sensor domain/sidechains that sense the electrical field across the membrane.

However, the predicted topology of Grs does not contain a voltage sensor domain like the well-described Shaker-type voltage-gated potassium channels [29]. In 2016, Barchad-Avitzur et al. showed that the agonist binding affinity of the GPCR M2 muscarinic acetylcholine receptor (M2R) is modulated by voltage, just like the EC_50_ values of AmGr3 (Fig. 2E) [30]. Using biophysical techniques in combination with site-directed mutagenesis, the authors identified a non-canonical tyrosine-based voltage sensor that appeared crucial for the voltage dependence of agonist binding to the M2R receptor. Whether the voltage dependence of AmGr3 is also based on tyrosine residues within the electrical field of the membrane and whether the voltage dependence of the fructose receptor plays a crucial physiological function for the perception of sugar concentrations honeybees remains to be shown.

However, it is important to verify the heterologous expression in the original organism to define the function of a receptor. In the fruit fly *Drosophila melanogaster,* this is often done via knock out or knock in mutants. In honeybees, there are no transposons available and RNAi works to a limited extend in nerve tissue, which makes the new CRIPR/Cas9 technique a very promising method for such scientific questions. Our study is the first to demonstrate that CRISPR/Cas9 is a successful method to investigate the function of taste receptors in adult honeybees on a behavioural level. Our results show that AmGr3 is a specialized fructose receptor in the honeybee, from both the biophysical characterization in oocytes and the behavioural perspective tested in honeybees. Healthy honeybees recognize both sucrose and fructose and respond more readily to increasing concentrations [31]. In our experiment, double nonsense mutations of the AmGr3 receptor led to a strong inhibition of responses to fructose, while responses to sucrose remained unaffected (Fig. 4, statistics in Tab. 2). Intriguingly, some bees with double nonsense mutations still responded to fructose. We cannot exclude the possibility that the sugar receptor AmGr1 and its co-receptor AmGr2 perceive fructose in a reduced manner when co-expressed in the same gustatory neuron (for sugar taste in *Drosophila melanogaster* a co-expression of multiple Grs is assumed to be necessary [25] [26]), although these receptors normally do not respond to fructose. Furthermore, other receptors in the antennae may react to water, the tactile stimuli or the osmolarity of the testing solution and thus generate the baseline measured for fructose. Nevertheless, AmGr1 and AmGr2 did not show any reaction towards fructose when tested in *Xenopus* oocytes [5]. Alternatively, one or several of the uncharacterized honeybee gustatory receptors might be able to perceive fructose, possibly through perceiving the molarity of liquids *per se*. In addition, fructose might be structurally similar to ligands of other gustatory receptors. Further characterization and investigation of the other taste receptors of the honeybee will bring clarity to these questions in the future.

## Conclusion

Our experiments demonstrate that CRISPR/Cas9 is an efficient tool to characterize taste receptors and other behaviourally relevant proteins in the honeybee. With the advent of this genetic tool, the honeybee has now a high potential for genetic manipulation. Taken together with the rich behavioural repertoire of this insect and its unique behavioural characteristics like division of labour, learning ability and dance language [1], this makes the honeybee an ideal model organism for studying gene function in a live insect. Furthermore, our data demonstrate that the AmGr3 receptor is not essential for larval development and that it is a specific fructose receptor in honeybee workers.

## Methods

To characterize the putative fructose receptor from *Apis mellifera*, we cloned the respective cDNA and expressed AmGr3 heterologously in *Xenopus laevis* oocytes. To elucidate the sugar perception and cation transport characteristics of AmGr3, its functional analysis was performed using the two-electrode voltage-clamp technique (TEVC). The electrical characteristics of AmGr3 were studied with respect to its sugar specificity, fructose affinity, cation selectivity and voltage dependency.

Confirming its function as a fructose receptor *in vivo* we used CRISPR/Cas9 in honeybee eggs [17] [16]. Mutated honeybees were raised in the laboratory [32]. At one-week of age, these animals were tested for their response to fructose and sucrose [21] [19]. The success of the mutation was controlled by fluorescence length analysis (FLA [33]) a next generation sequencing (NGS) [34] [35].

### *Xenopus* oocyte preparation

Investigations on AmGr3 were performed in oocytes of the African clawed frog *Xenopus laevis*. Permission for keeping *Xenopus* exists at the Julius-von-Sachs Institute and is registered at the government of Lower Franconia (reference number 70/14 and 55.2-2532-2-1035). Mature female *Xenopus laevis* frogs (healthy, non-immunized and not involved in any previous procedures) were kept at 20 °C at a 12/12 h day/night cycle in dark grey 96 l tanks (5 frogs/tank). Frogs were fed twice a week with floating trout food (Fisch-FitMast 45/7 2 mm, Interquell GmbH, Wehringen, Germany). Tanks are equipped with 30 cm long PVC pipes with a diameter of around 10 cm. These pipes are used as hiding places for the frogs. The water is continuously circulated and filtered by a small aquarium pump. For oocyte isolation, mature female *X. laevis* frogs were anesthetized by immersion in water containing 0.1 % 3-aminobenzoic acid ethylester. Following partial ovariectomy, oocytes were treated with collagenase I in Ca^2+^-free ND96 buffer (10 mM HEPES pH 7.4, 96 mM NaCl, 2 mM KCl, 1 mM MgCl_2_,) for 1 to 1.5 h. Subsequently, oocytes were washed with Ca^2+^-free ND96 buffer and kept at 16°C in ND96 solution (10 mM HEPESpH7.4, 96 mM NaCl, 2 mM KCl, 1 mM MgCl2, 1 mM CaCl2) containing 50 mg/l gentamycin. For electrophysiological experiments 10 ng of AmGr3 cRNA was injected into each stage V or VI oocyte. Oocytes were incubated for 2 to 3 days at 16 °C in ND96 solution containing gentamycin.

### RNA extraction and cDNA synthesis

For RNA extraction, frozen honeybee antennae, mouthparts and tarsi were broken up in 750 μl TriFast (peqGOLD, VWR, Radnor, USA) in a 2 ml Eppendorf (Hamburg, Germany) tube using Stainless Steel Beads (5 mm) and the TissueLyzer (QIAGEN, Venlo, Netherlands). After an incubation time of 5 min, 200 μl chloroform were added, mixed, centrifuged and the aqueous phase was applied to a PerfectBind RNA Colum of the Total RNA Kit (peqGOLD, VWR, Radnor, USA). Further extraction of total RNA was performed according to the kits protocol. RNA was precipitated with 3 M sodium acetate, washed with ethanol, dried and the pellet was resolved to adjust the concentration. Synthesis of cDNA was carried out with the AccuScript Hi-Fi cDNA Synthesis Kit (Agilent Technologies, Santa Clara, USA) using Oligo(dT) primer (18mers) according to the manufactures instructions and required concentrations. RNA was digested enzymatically with RNAse H (NEB, Ipswich, USA) following the protocol. According to the instructions, a large scale Phusion PCR (NEB, Ipswich, USA) was performed using a forward (5’-GAATTGTCTCGTTCGCAAATAC-3’) and a reverse primer (5’-CCGCTATTTACGAAAATTGG- 3’) covering the predicted ORF (open reading frame) of the *AmGr3* gene (NCBI: XM_016913387.1). The PCR product was applied and run on a 1 % (*w/v*) agarose gel. The appropriate band (1595 bp) was excised and purified as recommended by the Wizard SV Gel and PCR Clean-Up System (Promega, Fitchburg, USA). The blunt end PCR product was A-tailed with a 20 min incubation step at 72 °C by adding 0.2 mM dATP, taq polymerase and its required buffer (NEB, Ipswich, USA).

### Cloning and cRNA synthesis

Via T/A ligation the fragment was inserted in the pGEM-T vector following the manufactures recommendations (Promega, Fitchburg, USA). Competent *Escherichia coli* cells (*E. coli* JM109; Promega, Fitchburg, USA) were incubated with the ligation mixture on ice for 30 min and then transformed by a 45 sec heat shock at 42 °C. After cooling on ice, the cells could regenerate on the shaker (300 rpm) at 37 °C for 45 min in 500 μl LB medium (Carl Roth, Karlsruhe, Germany). They were subsequently plated on agar plates (LB agar; Carl Roth, Karlsruhe, Germany) containing Carbenicillin (100μg/ml; Carl Roth, Karlsruhe, Germany) and IPTG (1 M, 2.5 μl per plate; Carl Roth, Karlsruhe, Germany) and X-Gal (240 mM, diluted in Dimethylformamide, 37.5 μl per plate; Carl Roth, Karlsruhe, Germany) and could grow over night at 37 °C. Using blue-white selection, clones were picked, cultivated in a liquid overnight culture (LB and 100 μg/ml Carbenicillin; Carl Roth, Karlsruhe, Germany), pelleted and purified by the Plasmid Miniprep Kit I (peqGOLD, VWR, Radnor, USA). Inserts of the isolated plasmids were verified by sequencing. The complementary DNA (cDNA) of AmGr3 was then sub-cloned into oocyte expression vector pNBIu (based on pGEM vectors) by an advanced uracil-excision-based cloning technique using PfuX7 polymerase, as described by Nour-Eldin et al. (2006) and Nørholm (2010) [36] [37]. All constructs were verified by sequencing. For functional analysis, complementary RNA (cRNA) was prepared with the AmpliCap-Max T7 High Yield Message Maker Kit (Cellscript, Madison, WI, USA) according to the manufacturer’s specifications.

### Oocyte recordings

Solutions: In two-electrode voltage-clamp studies, oocytes were perfused with Tris/Mes-based buffers. The standard solutions contained 30 mM NaCl, 10 mM Tris/Mes (pH 7.4), 1 mM CaCl_2_, 1 mM MgCl_2_, and either 160 mM D-sorbitol (control solution) or 160 mM fructose. Solutions for cation selectivity measurements based on the standard solutions where NaCl was replaced by either 30 mM LiCl, KCl, RbCL or CsCl. For sugar specificity measurements, D-Sorbitol was exchanged by either 160 mM of fructose, glucose, sucrose, mannose, galactose, maltose, arabinose, raffinose, trehalose or melezitose. Osmolarity was adjusted to 220 mOsmol/L with D-sorbitol. For the determination of the fructose affinity of AmGr3, the fructose concentration in the standard solution varied between 0 and 500 mM. To balance the osmolarity, we compensated changes in the fructose concentration with D-sorbitol. Due to the high sugar concentration during the fructose dose-response measurements, the osmolarity was around 560 mOsmol/L, which was tolerated by the oocytes.

Electrical recordings and data analysis: For steady-state current (I_SS_) recordings with AmGr3 expressing oocytes, the standard voltage protocol was as follows: Starting from a holding potential (V_H_) of 0 mV, single 200 ms voltage pulses were applied from +40 to −150 mV in 20 mV decrements, unless otherwise stated in the figure legend. Fructose-induced currents were derived by subtracting the currents in the absence of fructose from the currents in its presence. For the calculation of the relative cation permeability of AmGr3, reversal potentials (V_rev_) were determined with either 30 mM KCl, LiCl, NaCl, RbCL or CsCl in the presence of 160 mM fructose. The relative permeability was calculated using the following equation [38]: 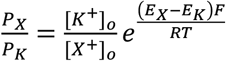, where [K^+^]_o_ is the external potassium concentration and [X^+^]_o_ is the external concentration of the test cation. E_K_ is the reversal potential with potassium and E_X_ is the reversal potential for the external test cation. F and R are the Faraday and gas constants, respectively, and T is the absolute temperature. For the calculation of EC_50_ values, the fructose dose-response curves at different membrane potentials were fitted with a hill equation: 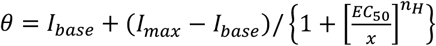, where I_base_ is the current in the absence of fructose, I_max_ the current in the presence of saturating fructose concentrations, EC_50_ the ligand concentration where the half maximal activity of AmGr3 is reached, x is the ligand concentration and nH is the Hill-coefficient.

### Preparation of sgRNA

Appropriate sites for sgRNAs (single guide RNA) were found in the first exons of the ORF (open reading frame) of the putative fructose receptor *AmGr3* in the genome of *Apis mellifera*. Using benchling (https://benchling.com, San Francisco, USA) we defined the target specific crRNA to be 20bp long, next to an NGG pam site and to start with a guanine base (for position within the gene also see Fig. 3). A sequence with a minimal on-target score of 50 % and an off-target score of at least 97 % were chosen. The secondary structure of the whole sgRNA was tested with the Vienna sgRNA fold program (http://rna.tbi.univie.ac.at/cgi-bin/RNAWebSuite/RNAfold.cgi, University of Wien, Austria) to assure that its stable part (tracrRNA) folds into the interaction structure for the Cas9 enzyme and the 20 bp of the variable part (crRNA) is still freely accessible and can thus bind the genomic target. Two primers with overlapping sequences were designed. The forward primer was containing a T7 promoter and the certain crRNA sequence (5’-GAAATTAATACGACTCACTATA-**GCAACTTGTAGTGATGTGCT-**<u>GTTTTAGAGCTAGAAATAGC</u>-3’), the reverse primer was containing the tracrRNA sequence (5’-AAAAGCACCGACTCGGTGCCACTTTTTCAAGTTGATAACGGACTAGCCTTATTTTAACTT-<u>GCTATTTCTAGCTCTAAAAC</u>-3’). Both were processed by an overlapping Phusion PCR (NEB, Ipswich, USA) and purified with Monarch PCR & DNA Cleanup Kit (5 μg) (NEB, Ipswich, USA), checked on an 1 % (*w/v*) agarose gel and quantified (NanoDrop BioPhotometer plus; Eppendorf, Hamburg, Germany). The PCR product was the template for the sgRNA synthesis according to the protocol of the RiboMAX Large Scale RNA Production Systems with T7 RNA polymerase (Promega, Fitchburg, USA). After the kits DNAse digestion, sgRNA was purified with MEGAclear Transcrition Clean-Up Kit (Invitrogen, Carlsbad, USA), checked on an agarose gel and quantified (NanoDrop BioPhotometer plus; Eppendorf, Hamburg, Germany). The sgRNA was aliquoted and frozen in portions. Initially, we produced three different sgRNAs and tested them in different concentrations. In this preliminary experiment (data not shown), we defined the hatching and mutation rates for each sgRNA and their best ratio with Cas9 enzyme. During the experiment a fresh aliquot with 46 ng/μl sgRNA and 3.13 μM commercial Cas9 enzyme (Cas9 Nuclease, S. pyogenes, 20 μM; NEB, Ipswich, USA) was used for each day and stored on ice.

### Honeybee egg harvest

Nine hives with related and naturally inseminated queens of *Apis mellifera carnica* were kept at the bee station of the Julius-Maximilians-University of Würzburg in July and August 2018. Bees were allowed to forage freely. In order to stimulate the oviposition of the queen, the colonies were fed with ApiInvert or ApiFonda (Südzucker, Mannheim, Germany) during bad weather or insufficient floral nectar flow. For egg harvest the queens were locked in the JENTER system (Karl-Heinz Jenter, Frickenhausen, Germany) the evening before. As a result, they were forced to lay their eggs through a comb-like cell grid and onto removable JENTER plug-in cells. All plug-in cells were placed on prefabricated plates so that they could be easily exchanged at once. The queens were left in the system for three days and the overnight eggs were discarded. For injection, eggs were harvested every 1.5 – 2 h, starting in the morning of each day. The sum of the injection time and the time since the last harvest was set for 3 h each round. Thus, the eggs were not older than 3 hours and should not have gone through any maturity division when injected. For the transport of the eggs we used an isolated transport box with preheated packs (35 °C, kept in the climate chamber).

### Microinjection of eggs

The eggs were processed and injected in a climate chamber maintained at constant 35 °C with no humidity regulation. For this purpose, the egg-containing plug-in cells were removed from the plates and fixed vertically on petri dishes (VWR) with plastiline (Pelikan Schindellegi, Switzerland). Thus, the tops of the eggs were easily accessible on the outer ring, while the eggs were attached to the cell at their bottom. The injection area was surrounded by a box with a glass lid and a liquid reservoir to ensure humidity during the injection process. In this area, the rings could be rotated with one hand while the injection needle entered it through a small hole. The ICSI Glass Pipettes used (BioMedical Instruments, Zöllnitz, Germany) were controlled with the Singer Mk1 micromanipulator (SINGER instruments, Somerset, England) and inserted into the upper quarter of the eggs. For injection, the PLI-100A picolitre injector (Warner Instruments, Hamden, USA) with a footswitch was used and operated by the climate chamber’s air system (max. 7 bar, with intermediate filter). Each egg was injected with 400 pl of either water or sgRNA6 with Cas9 (prepared as described above; injection time: 120 ms; P_bal_: 5 kPa; P_inje_: 60 kPa). Thereafter, the rings with injected eggs were placed in plastic boxes with a sulphurous atmosphere (1ml of 16 % sulfuric acid per liter of volume, separated from the rings by a grid) to keep the puncture site sterile. One day after the injection, burst or dried eggs were removed. The 24 h mortality rate reflects the failure of the egg to develop due to injection, because it is approximately equal for eggs injected with water or sgRNA and Cas9 enzymes. A few hours before hatching the sulfuric acid was washed off well and replaced with water. Immediately after hatching, the larvae were removed carefully with a modified Chinese grafting tool. The hatching rate displays the tolerance of the sgRNA with Cas9 enzyme, since it is over 95 % when the eggs were injected with water only. In our experiment, we performed two replicates of injection weeks.

### Artificial rearing of honeybees

The freshly hatched honeybee larvae were carefully detached from the plug-in cells with a modified Chinese grafting tool dipped in larvae food. They were placed laterally in prepared Nicot-wells (NICOTPLAST, Maisod, France) containing larval food. Care was taken not to contaminate the lateral breathing holes of the upper side. The food and rearing procedures are described in detail in Schmehl et al. (2016, [32]) with some deviations from the protocol). The Nicot-wells in which the larvae were placed were already filled with “larval food A and B”. They were placed in 48-well NUNC plates (ThermoFisher, Massachusetts, USA) in which they lay on cotton wool slices soaked with 0,4 % MBC (methylbenzethoniumchlorid chloride, *w/v*) and glycerol (84.5 % and 15.5 %, *v/v*). The closed 48-well plates rested in a separate box in the incubator at 35°C for the duration of larval development. As described in Schmehl et al. (2016, [32]), the box contained a K_2_SO_4_ buffer which adjusted the humidity to ~ 94 %. After the larvae consumed the food of all conducted feedings (for feeding ingredients and times see also [32]), they were transferred to sterile filter paper in a fresh 48-well plate. During pupation, the animals were left to develop in a ~ 75 % humidity, which was achieved by NaCl buffer. After hatching within the 48-well plates, the adult bees were individually marked with colored number plates (Opalith queenmarking plates) using super glue (UHU). After cutting of a wing for easier handling and safety reasons, they were placed in a cage with pollen and sugar water (20 % sucrose, 10 % fructose and 10 % glucose, *w/w/w/v*). All bees of one replicate shared one cage, including the labelled control animals, and were kept in an incubator maintained at 28 °C.

### Testing responsiveness to sucrose and fructose

Bees were tested for their proboscis extension response (PER) to increasing concentrations of sucrose and fructose at one-week old. For this test, each bee was immobilized on ice, carefully mounted in brass tubes and fixed with adhesive tape [19]. At each test, both antennae were stimulated with a droplet of a certain sugar water concentration. Both sugars, alternatingly starting with fructose or sucrose, were tested. After a test with water, the test of a sugar solution was carried out with the following increasing concentrations 16 %, 20 %, 25 %, 32 %, 40 %, 50 % and 63 % (*w/v*) which corresponds to a logarithmic series of approximately 1.2; 1.3; 1.4; 1.5; 1.6; 1.7; 1.8. Contaminations occurring at the antennae were immediately removed and rinsed with water. It was already shown that sucrose responsiveness is not affected by the order of the concentrations tested [19]. The positive PER for each concentration was recorded individually for each sugar (sucrose or fructose) and each bee. To prevent intrinsic sensitization, there was an inter-trial interval of 2 min [19]. The sum of the responses to water and the ascending concentrations of the certain sugar displays the gustatory response score (GRS) of a bee for fructose or sucrose.

### Genotyping via fluorescence length analysis and next generation sequencing

Directly after the behavioral test, the bees were individually immersed in liquid nitrogen and stored at −20 °C. Their heads were dissected, placed in 2ml Eppendorf (Hamburg, Germany) tubes and disrupted with a pre-cooled Stainless-Steel Beads (5 mm) in the TissueLyzer (QIAGEN, Venlo, Netherlands). The genomic DNA (gDNA) of each bee was relieved by 200 μl CTAB lysis buffer (1 % CTAB (*w/v*), 50 mM Tris pH8, 10 mM EDTA, 0.75 M NaCl) and 2 μl protein kinase K (NEB, Ipswich, USA) during 2 h at 60 °C. It was isolated by phenol/chloroform/isoamyl alcohol (25:24:1, pH 7.5-8.0), washed with chloroform (250 μl) and precipitated with natrium acetate (3 M; 20μl) and ice-cold ethanol (100 %; 450 μl). After washing with 70 % ethanol, final centrifuging and drying, the pellets were resolved in nuclease-free water (100 μl each). With the obtained gDNA samples, a PCR was performed with a hex-labeled forward primer (5’-HEX-TGCGTACTTGTATTACTACTTAGTGC-3’) and a reverse primer (5’-AACAAGTTGCAAATATTTCCAACGG-3’), both framing the sgRNA site. In 96-well quality PCR plates (for FLA, Kisker Biotech, Steinfurt, Germany) 1 μl of each PCR product was edited with Hi-Di Formamide (20 μl, ThermoFisher, Massachusetts, USA) and Gene Scan 500 ROX dye Size Standard (0.5 μl, ThermoFisher, Massachusetts, USA) and examined in a fluorescence length analysis via the HEX-label. The obtained peaks accurately display length deviations of only 1 bp from the wildtype (evaluated with PeakScanner2; ThermoFisher, Massachusetts, USA). To ensure that these shifted peaks represent mutations in the genomic DNA we performed next generation sequencing (NGS, performed with GENEWIZ, Leipzig, Germany) with all candidate samples. Samples were first indexed with two tags (5’-CTGTGATG-3’ and 5’-GCGCAATA-3’) for multiplexing and amplified with adapter overhangs (complete sequences, forward: 5’-ACACTCTTTCCCTACACGACGCTCTTCCGATCTCTGTGATGtgcgtacttgtattactacttagtg-3’ and reverse: 5’- ACACTCTTTCCCTACACGACGCTCTTCCGATCTGCGCAATAtgcgtacttgtattactacttagtg-3’) for a second multiplexing process to be performed at GENEWIZ directly before sequencing on a Illumina HiSeq 2500 (2×250bp, Rapid Run). We demultiplexed the samples by the barcoding using HMMer v3.2.1 [39]. Forward and reverse reads were merged and subsequently quality filtered (maxEE=1, minlen=100) using USEARCH v11 [40]. We then identified indel lengths and counted variants with an own perl script for each sample. Since each animal has two alleles, we classified each of them with “wt” (for wildtype) if in-frame indels were a multiple of 3 bps, leaving the open reading frame intact. Nonsense alleles were labeled with “ns”, including open reading frame shifts and leading to non-functional proteins (see Fig. 3). Worker bees may be homozygous or heterozygous combining these possible configurations (derived from two chromosomes). For the investigation of the behavior only homozygous wildtypes (wt/wt) and mutants (ns/ns) were analyzed.

### Quantification and statistical analysis

All electrophysiological experiments were performed at least twice (independent experiments with oocytes from different batches). Sample size, n, and statistical details (mean ± standard error, SE or standard deviation, SD) are given in the figure legends for each experiment. For statistical analysis, the software Igor Pro 8 (waveMetrics, Inc., Lake Oswego, Oregon, USA) and Excel (Microsoft Corp. Redmond, Washington, USA) was used.

For structural prediction of the AmGr3 protein (Fig. 3) the sequence was modelled to the Cryo-EM structure of *Apocrypta bakeri* Orco (PDB 6c70A, [27]) using I-Tasser (University of Michigan, [41], [42]) and compared with other predictions (PHYRE2, Imperial College London and TMHMM, DTU Bioinformatics Denmark).

The GraphPad Prism software (version 7.03; GraphPad Software, San Diego, USA) was used for analyzing survival and hatching. Fisher’s exact tests were used to compare 24h survival and hatching of eggs in both replicates either injected with sgRNA and Cas9 or with water. Chi-Square tests was applied to compare these in total values additionally including not injected eggs. The fructose and sucrose gustatory response curves were analyzed with the IBM SPSS software (version 23.0.0.0; IBM, New York, USA) via logistic regression [factor genotype] and graphical displayed in Graph Pad Prism.

## Supplementary information

### Additional files

**Xlsx-file 1:** suppl_24h_3d_survival_injection; **Xlsx-file 2:** suppl_FLA_vs_NGS_raw_scripted_reads, **Xlsx-file 3:** suppl_PER_data_FRUC_SUC; **Xlsx-file 4:** suppl_portion_wt_ns_FLA_peaks; **pl-file 1:** suppl_SCRIPT_get_indels.

## Acknowledgments

We thank Anne Huber and Sandra Reimlinger for biophysical analysis of AmGr3 in oocytes, Marianne Otte and Vivien Bauer for help and assistance in CRISPR/Cas9 related molecular work and injection procedure. We thank Markus Thamm for molecular lab expertise and Karin Möller for technical assistance. We thank Maria Scherrer, Lisa Rauscher, Lioba Hilsmann, Tabea Lammert, Daniel Rodriquez, Katharina Beer and Felix Schilcher for honeybee queen caging, egg collection, preparation for injection and help with artificial rearing. We thank Martin Gabel, Gaby Läbisch and Dirk Ahrens for the beehive maintenance. This project was supported by a fund of the DFG to R.S. (SCHE 1573/8-1).

## Author Contributions

D.G., L.D., M.B., A.K. and R.S. designed research and wrote the paper. I. S.-D. devised experiments and contributed to manuscript revision. D.G. and F.L.R.F. performed biophysical characterization experiments and analysed the data. L.D. performed CRISPR/Cas9 experiments and data analysis. B.K. was involved in artificial rearing of honeybees. AK performed bioinformatics for genotyping.

## Fundings

This study was supported by the Deutsche Forschungsgemeinschaft through grant SCHE 1573/8-1 to R.S.

## Availability of data and materials

Correspondence and requests for materials not included in the supplementary information should be addressed to L.D. and M.B. for CRISPR/Cas9, to L.D. for *in vitro* rearing of larvae, to D.G. and F.L.R.F. for characterization and to A.K. for bioinformatics.

## Ethics approval and consent to participate

No ethics approval or consent to participate was required for this study.

## Competing interests

The authors declare that they have no competing interests.

## Author details

1 University of Würzburg, Behavioral Physiology and Sociobiology, Biocenter, Am Hubland, 97074 Würzburg, Germany, 2 University of Würzburg, Julius-von-Sachs-Institute, Molecular Plant Physiology and Biophysics, Biocenter, Julius-von-Sachs-Platz 2, 97082 Würzburg, Germany, 3 University of Würzburg, Department of Bioinformatics, Biocenter, Am Hubland, 97074 Würzburg, Germany, 4 University of Würzburg, Animal Ecology and Tropical Biology, Biocenter, Am Hubland, 97074 Würzburg, Germany, 5 Heinrich-Heine-Universität Düsseldorf, Evolutionary Genetics, Universitätsstraße 1,40225 Düsseldorf, Germany.

## References

[1] T. D. Seeley, The wisdom of the hive: the social physiology of honey bee colonies, Harvard University Press, 2009.

[2] J. M. Graham, The Hive and the Honeybee, Hamilton, Illinois: Dadant & Sons, 1992.

[3] M. Bertazzini and G. Forlani, “Intraspecific variability of floral nectar volume and composition in rapeseed (Brassica napus L. var. oleifera),” Frontiers in plant science, no. 7, p. 288, 2016.

[4] Robertson, H. M. and Wanner, K. W., “The chemoreceptor superfamily in the honey bee, Apis mellifera: expansion of the odorant, but not gustatory, receptor family,” Genome research, vol. 16, no. 11, pp. 1395–1403, 2006.

[5] Jung, J. W., Park, K. W., Ahn, Y. J. and Kwon, H. W, “Functional characterization of sugar receptors in the western honeybee, Apis mellifera,” Journal of Asia-Pacific Entomology, vol. 18, no. 1, pp. 19–26, 2015.

[6] Takada, T., Sasaki, T., Sato, R., Kikuta, S. and Inoue, M. N., “Differential expression of a fructose receptor gene in honey bee workers according to age and behavioral role,” Archives of insect biochemistry and physiology, vol. 97, no. 2, p. e21437, 2018.

[7] S. S. Haupt, “Central gustatory projections and side-specificity of operant antennal muscle conditioning in the honeybee,” Journal of Comparative Physiology A, vol. 193, no. 5, pp. 523–535, 2007.

[8] Simcock, N. K., Wakeling, L. A., Ford, D. and Wright, G. A., “Effects of age and nutritional state on the expression of gustatory receptors in the honeybee (Apis mellifera),” PloS one, vol. 12, no. 4, p. e0175158, 2017.

[9] D. A. Stanley, D. Gunning and J. C. Stout, “Pollinators and pollination of oilseed rape crops (Brassica napus L.) in Ireland: ecological and economic incentives for pollinator conservation,” Journal of Insect Conservation, vol. 17, no. 6, pp. 1181–1189, 2013.

[10] V. R. Chalcoff, M. A. Aizen and L. Galetto, “Nectar concentration and composition of 26 species from the temperate forest of South America,” Annals of botany, vol. 97, no. 3, pp. 413–421, 2006.

[11] Sato, K, Tanaka, K. and Touhara, K., “Sugar-regulated cation channel formed by an insect gustatory receptor,” Proceedings of the National Academy of Sciences, vol. 108, no. 28, pp. 11680–11685, 2011.

[12] Yoo, S. D., Cho, Y. H. and Sheen, J., “Arabidopsis mesophyll protoplasts: a versatile cell system for transient gene expression analysis,” Nature protocols, vol. 2, no. 7, p. 1565, 2007.

[13] S. C. Lin, Y. Y. Chang and C. C. Chan, “Strategies for gene disruption in Drosophila,” Cell & bioscience, vol. 4, no. 1, p. 63, 2014.

[14] Bassett, A. R., Tibbit, C., Ponting, C. P. and Liu, J. L., “Highly efficient targeted mutagenesis of Drosophila with the CRISPR/Cas9 system,” Cell reports, vol. 4, no. 1, pp. 220–228, 2013.

[15] Basu, S., Aryan, A., Overcash, J.M., Samuel, G.H., Anderson, M.A., Dahlem, T.J., Myles, K.M. and Adelman, Z.N., “Silencing of end-joining repair for efficient site-specific gene insertion after TALEN/CRISPR mutagenesis in Aedes aegypti,” Proceedings of the National Academy of Sciences, vol. 112, no. 13, pp. 4038–4043, 2015.

[16] Kohno, H., Suenami, S., Takeuchi, H., Sasaki, T. and Kubo, T., “Production of Knockout Mutants by CRISPR/Cas9 in the European Honeybee, Apis mellifera L.,” Zoological science, vol. 33, no. 5, pp. 505–513, 2016.

[17] Roth, A., Vleurinck, C., Netschitailo, O., Bauer, V., Otte, M., Kaftanoglu, O., Page, R.E. and Beye, M., “A genetic switch for worker nutrition-mediated traits in honeybees,” PLoS biology, vol. 17, no. 3, p. e3000171, 2019.

[18] S. V. Prykhozhij, S. Steele, B. Razaghi and J. N. Berman, “A rapid and effective method for screening, sequencing and reporter verification of engineered frameshift mutations in zebrafish,” Disease models & mechanisms, vol. 10, no. 6, pp. 811–822, 2017.

[19] Scheiner, R., Abramson, C.I., Brodschneider, R., Crailsheim, K., Farina, W.M., Fuchs, S., Gruenewald, B., Hahshold, S., Karrer, M., Koeniger, G. and Koeniger, N., “Standard methods for behavioural studies of Apis mellifera,” Journal of Apicultural Research, vol. 52, no. 4, pp. 1–58, 2013.

[20] Scheiner, R., Page Jr, R. E. and Erber, J., “The effects of genotype, foraging role, and sucrose responsiveness on the tactile learning performance of honey bees (Apis mellifera L.),” Neurobiology of learning and memory, vol. 76, no. 2, pp. 138–150, 2001.

[21] Scheiner, R., Page, R. E. and Erber, J., “Sucrose responsiveness and behavioral plasticity in honey bees (Apis mellifera),” Apidologie, vol. 35, no. 2, pp. 133–142, 2004.

[22] Wright, G.A., Choudhary, A.F. and Bentley, M.A., “Reward quality influences the development of learned olfactory biases in honeybees,” Proceedings of the Royal Society B: Biological Sciences, vol. 276, no. 1667, pp. 2597–2604, 2009.

[23] de Brito Sanchez, G., Ortigao-Farias, J.R., Gauthier, M., Liu, F. and Giurfa, M., “Taste perception in honeybees: just a taste of honey?,” Arthropod-Plant Interactions, vol. 1, no. 2, pp. 69–76, 2007.

[24] Miyamoto, T., Slone, J., Song, X. and Amrein, H., “A fructose receptor functions as a nutrient sensor in the Drosophila brain,” Cell, vol. 151, no. 5, pp. 1113–1125, 2012.

[25] Jiao, Y., Moon, S. J., Wang, X., Ren, Q. and Montell, C., “Gr64f is required in combination with other gustatory receptors for sugar detection in Drosophila,” Current Biology, vol. 18, no. 22, pp. 1797–1801, 2008.

[26] Slone, J., Daniels, J. and Amrein, H., “Sugar receptors in Drosophila,” Current Biology, vol. 17, no. 20, pp. 1809–1816, 2007.

[27] Butterwick, J. A., del Mármol, J., Kim, K. H., Kahlson, M. A., Rogow, J. A., Walz, T. and Ruta, V., “Cryo-EM structure of the insect olfactory receptor Orco,” Nature, vol. 560, no. 7719, p. 447, 2018.

[28] Robertson, H. M., “Molecular evolution of the major arthropod chemoreceptor gene families,” Annual review of entomology, vol. 64, pp. 227–242, 2019.

[29] F. Bezanilla, “The voltage-sensor structure in a voltage-gated channel,” Trends in biochemical sciences, vol. 30, no. 4, pp. 166–168, 2005.

[30] Barchad-Avitzur, O., Priest, M. F., Dekel, N., Bezanilla, F., Parnas, H. and Ben-Chaim, Y., “A novel voltage sensor in the orthosteric binding site of the M2 muscarinic receptor,” Biophysical journal, vol. 111, no. 7, pp. 1396–1408, 2016.

[31] Değirmenci, L., Thamm, M. and Scheiner, R., “Responses to sugar and sugar receptor gene expression in different social roles of the honeybee (Apis mellifera),” Journal of insect physiology, vol. 106, pp. 65–70, 2018.

[32] Schmehl, D. R., Tomé, H. V., Mortensen, A. N., Martins, G. F. and Ellis, J. D., “Protocol for the in vitro rearing of honey bee (Apis mellifera L.) workers,” ournal of Apicultural Research, vol. 55, no. 2, pp. 113–129, 2016.

[33] M. K. Ramlee, T. Yan, A. M. Cheung, C. T. Chuah and S. Li, “High-throughput genotyping of CRISPR/Cas9-mediated mutants using fluorescent PCR-capillary gel electrophoresis,” Scientific reports, vol. 5, no. 1, pp. 1–13, 2015.

[34] J. Shendure and H. Ji, “Next-generation DNA sequencing,” Nature biotechnology, vol. 26, no. 10, p. 1135, 2008.

[35] B. F. Vogel, V. Fussing, B. Ojeniyi, L. Gram and P. Ahrens, “High-resolution genotyping of Listeria monocytogenes by fluorescent amplified fragment length polymorphism analysis compared to pulsed-field gel electrophoresis, random amplified polymorphic DNA analysis, ribotyping, and PCR–restriction fragment length po,” Journal of food protection, vol. 67, no. 8, pp. 1656–1665, 2004.

[36] H. H. Nour-Eldin, B. G. Hansen, M. H. Nørholm, J. K. Jensen and B. A. Halkier, “Advancing uracil-excision based cloning towards an ideal technique for cloning PCR fragments,” Nucleic acids research, vol. 34, no. 18, pp. e122–e122, 2006.

[37] M. H. Nørholm, “A mutant Pfu DNA polymerase designed for advanced uracil-excision DNA engineering,” BMC biotechnology, vol. 10, no. 1, p. 21, 2010.

[38] D. Becker, I. Dreyer, S. T. E. F. A. N. HoTH, J. D. Reid, H. Busch, M. Lehnen, K. Palme and R. Hedrich, “Changes in voltage activation, Cs+ sensitivity, and ion permeability in H5 mutants of the plant K+ channel KAT1,” Proceedings of the National Academy of Sciences, vol. 93, no. 15, pp. 8123–8128, 1996.

[39] S. R. Eddy, “Eddy, Sean R. “Accelerated profile HMM searches,” PLoS computational biology, vol. 10, no. 7, p. e1002195, 2011.

[40] R. C. Edgar, “Search and clustering orders of magnitude faster than BLAST,” Bioinformatics, vol. 19, no. 26, pp. 2460–2461, 2010.

[41] J. Yang and Y. Zhang, “I-TASSER server: new development for protein structure and function predictions,” Nucleic acids research, vol. 43, no. W1, pp. W174–W181, 2015.

[42] C. Zhang, P. L. Freddolino and Y. Zhang, “COFACTOR: improved protein function prediction by combining structure, sequence and protein–protein interaction information,” Nucleic acids research, vol. 45, no. W1, pp. W291–W299, 2017.

[43] M. Jinek, K. Chylinski, I. Fonfara, M. Hauer, J. A. Doudna and E. Charpentier, “A programmable dual-RNA–guided DNA endonuclease in adaptive bacterial immunity,” science, vol. 337, no. 6096, pp. 816–821, 2012.

[44] J. M. Graham, “The Hive and the Honeybee, Dadant & Sons,” Hamilton, Illinois, 1992.

[45] M. Bertazzini and G. Forlani, “Intraspecific variability of floral nectar volume and composition in rapeseed (Brassica napus L. var. oleifera),” Frontiers in plant science, vol. 7, p. 288, 2016.

